# Production of the infant formula ingredient 1,3-olein-2-palmitin in Arabidopsis seeds

**DOI:** 10.1101/2020.10.01.315424

**Authors:** Harrie van Erp, Fiona M Bryant, Jose Martin-Moreno, Peter J Eastmond

**Author notes:** **Correspondence:** Peter J. Eastmond, Rothamsted Research, Harpenden, Hertfordshire, AL5 2JQ, UK. **One-sentence summary:** Combined fatty acid and glycerolipid biosynthetic pathway engineering enables seeds to produce the infant formula ingredient 1,3-olein-2-palmitin.

## Abstract

In human milk fat, palmitic acid (16:0) is esterified to the middle (sn-2 or β) position on the glycerol backbone and oleic acid (18:1) predominantly to the outer positions, giving the triacylglycerol (TG) a distinctive stereoisomeric structure that is believed to assist nutrient absorption in the infant gut. However, the fat used in most infant formulas is derived from plants, which preferentially esterify 16:0 to the outer positions. We have previously showed that the metabolism of the model oilseed *Arabidopsis thaliana* can be engineered to incorporate 16:0 into the middle position of TG. However, the fatty acyl composition of Arabidopsis seed TG does not mimic human milk, which is rich in both 16:0 and 18:1 and is defined by the high abundance of the TG molecular species 1,3-olein-2-palmitin (OPO). Here we have constructed an Arabidopsis *fatty acid biosynthesis 1-1 fatty acid desaturase 2 fatty acid elongase 1* mutant with around 20% 16:0 and ~70% 18:1 in its seeds and we have engineered it to esterify more than 80% of the 16:0 to the middle position of TG, using heterologous expression of the human lysophosphatidic acid acyltransferase isoform AGPAT1, combined with suppression of LYSOPHOSPHATIDIC ACID ACYLTRANSFERASE 2 and PHOSPHATIDYLCHOLINE:DIACYLGLYCEROL CHOLINEPHOSPHOTRANSFERASE. Our data suggest that oilseeds can be engineered to produce TG that is rich in OPO, which is an important structured fat ingredient used in infant formulas.

## INTRODUCTION

Human milk is considered the optimal source of nutrition for infants and it is their main food during the first 4–6 months of life (Innis 2011; Wei et al., 2019). The lipid fraction provides approximately half the infant’s calories and mainly consists of triacylglycerols (TG), which account for about 98% of total lipids (Wei et al., 2019). Palmitic acid (16:0) is the most abundant saturated fatty acid (FA) in human milk, providing about 20–25% of the total milk FAs (Wei et al., 2019). In human milk, over 70% of this 16:0 is esterified to the middle (sn-2 or β) positon on the glycerol backbone of TG, while unsaturated FA, such as oleic acid (18:1), occupy the outer (sn-1/3 or α) positions (Breckenridge et al., 1969; Giuffrida et al., 2018). This allow greater efficiency of 16:0 absorption and utilization in infants when compared to infants fed with TG containing 16:0 preferentially esterified to the sn-1/3 positions (Innis 2011; Béghin et al., 2018). This is because during digestion, lipases release FAs preferentially from the sn-1/3 positions of TG to produce FA and 2-monoacylglycerols (Innis 2011; Béghin et al., 2018). 16:0 esterified to the sn-2 position of monoacyglycerol is easily absorbed while free 16:0 tends to form insoluble calcium soaps (Innis 2011; Béghin et al., 2018).

In infant formulas the lipid phase is usually provided by a mixture of vegetable fats and oils, blended to mimic the FA composition of human milk (Wei et al., 2019). However, plants virtually always exclude 16:0 from the sn-2 position of their TG (Brockerhoff and Yurkowsk, 1966; Christie et al., 1991) and therefore vegetable fats cannot mimic the regiospecific distribution of 16:0 that is found in human milk. To address this problem several companies have developed human milk fat substitutes (HMFS) that are produced by enzyme-catalyzed acidolysis (or alcoholysis and esterification) of fractionated vegetable fats and free FAs using sn-1/3-regioselective lipases (Wei et al., 2019). These HMFS contain TG with substantial levels of 16:0 enrichment at the sn-2 position and are widely use as ingredients in infant formulas (Wei et al., 2019). However, they are relatively costly to make, and it remains technically challenging to manufacture a true mimetic with over 70% of 16:0 esterified to the sn-2 position (Ferreira-Dias and Tecelão, 2014).

For this reason, we decided to investigate whether plant lipid metabolism can be engineered to directly produce TG with 16:0 enrichment at the sn-2 position (van Erp et al., 2019). TG is formed by a cytosolic glycerolipid biosynthetic pathway on the endoplasmic reticulum (ER) and the enzyme responsible for acylation of the sn-2 position is lysophosphatidic acid acyltransferase (LPAT) (Fig. 1) (Ohlrogge and Browse, 1995). ER-localized LPAT isoforms discriminate against a 16:0-Coenzyme A (CoA) substrate (Kim et al., 2005). By contrast, chloroplast-localized LPAT isoforms are highly selective for 16:0-acyl carrier protein (ACP) and will also accept 16:0-CoA (Joyard and Douce, 1977; Frentzen et al., 1983). We therefore modified a chloroplast LPAT by removing its targeting signal so that it localizes to the ER (ΔCTS-LPAT1 or mLPAT1) and expressed it in the model oilseed *Arabidopsis thaliana*, where it esterified >30% of 16:0 at the sn-2 position of TG (van Erp et al., 2019). Additionally, by suppressing endogenous ER-localized LPAT2 activity to relieve competition (van Erp et al., 2015) and by knocking out phosphatidylcholine:diacylglycerol cholinephosphotransferase (PDCT) to reduce diacylglycerol (DG) flux through phosphatidylcholine (PC) (Bates et al., 2012), we were able to increase the percentage of 16:0 at sn-2 to ~70% (van Erp et al., 2019).

**Figure 1.**
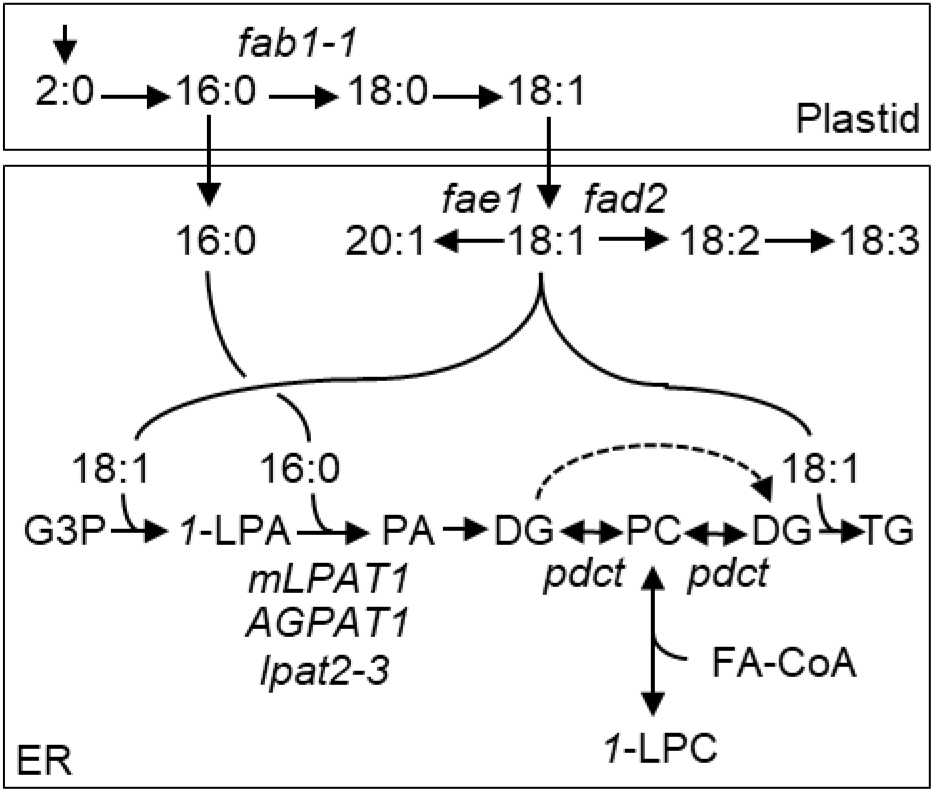
A simplified diagram illustrating the strategy used to produce OPO in Arabidopsis seeds. A combination of the hypomorphic *fab1-1* and null *fae1* and *fad2* mutant alleles was used to produce high levels of 16:0 and 18:1 in seeds. Expression of an ER-localised LPAT with 16:0-CoA preference combined with a hypomorphic *lpat2-3* and null *pdct* mutant allele was then used to enable 16:0 to be preferentially esterified to the sn-2 position of *1*-LPA and the products channelled into TG. 16:0, palmitic acid; 18:1, oleic acid; CoA, Coenzyme A; G3P, glycerol-3-phosphate; *1*-LPA, sn-1 lysophosphatidic acid; PA, phosphatidic acid, DG, diacylglycerol, TG, triacylglycerol; PC, phosphatidylcholine; *1*-LPC, sn-1 lysophosphatidylcholine; FA, fatty acid.

Although our work showed that plant lipid metabolism can be engineered to preferentially esterify 16:0 to the sn-2 position in TG (van Erp et al., 2019), the total FA composition of the Arabidopsis seeds does not resemble human milk and the TG we produced would not be appropriate for use as an infant formula ingredient. 16:0 is ~3-fold less abundant in Arabidopsis seeds and they contain a high proportion of polyunsaturated and very-long-chain FA species that are essentially absent from human milk. The most abundant FA in human milk is 18:1 and, because of the unusual regiospecific distribution of the next most abundant FA 16:0, the major molecular species of TG is usually 1,3-olein-2-palmitin (OPO) accounting for ~14% of the total (Giuffrida et al., 2018). The aim of this study was to investigate whether Arabidopsis seeds can be engineered to produce OPO, by combining 16:0 enrichment at the sn-2 position in TG with a total FA composition rich in the appropriate ratio of 16:0 and 18:1.

## RESULTS

### Seed of *fab1-1 fae1 fad2* are high in 16:0 and 18:1 (HPHO)

To obtain Arabidopsis seeds with 16:0 content equivalent to human milk, the level must be increase ~3-fold to 20-25%. One approach to achieve this is to reduce fatty acid synthase catalysed 16:0 elongation by disrupting the β-ketoacyl-ACP synthase II gene *FATTY ACID BIOSYNTHESIS 1* (*FAB1*) (Fig. 1) (Wu et al., 1994; Carlsson et al., 2002). *FAB1* is an essential gene in Arabidopsis (Pidkowich et al., 2007), but a single hypomorphic *fab1-1* allele has been characterised that contains ~17% 16:0 in its seeds (Wu et al., 1994; Carlsson et al., 2002). In *fab1-1*, 16:0 can then be increased to ~24% by disrupting FATTY ACID ELONGASE 1 (FAE1) (James and Dooner, 1991), which is required for very-long-chain fatty acid synthesis (Fig. 1) (James et al., 1995; Millar and Kunst, 1997). However, *fab1-1 fae1* seeds still contain a high proportion of polyunsaturated fatty acids (James and Dooner, 1991), produced via FATTY ACID DESATURASE 2 (FAD2) (Fig. 1) (Miquel and Browse, 1992; Okuley et al., 1994). To create a background high both 16:0 and 18:1 (HPHO) we therefore constructed a *fab1-1 fae1 fad2* mutant by crossing (James and Dooner, 1991). Analysis of homozygous *fab1-1 fae1 fad2* seeds showed that the FA composition of the TG is high in 16:0 and 18:1, which account for ~20 and ~70% for total FA, respectively (Fig. 2). Other FA species that are normally abundant in wild type Arabidopsis Col-0 seed TG, such as linoleic acid (18:2), linolenic acid (18:3) and eicosenoic acid (20:1), each account for <3% (Fig. 2). Comparison with the double mutants showed that 16:0 content in *fab1-1 fae1* is reduced significantly (P > 0.05) in the *fad2* background (Fig. 2). Nevertheless, the HPHO composition of *fab1-1 fae1 fad2* seeds suggests that this genetic background is appropriate to test whether OPO can be produced in seeds.

**Figure 2.**
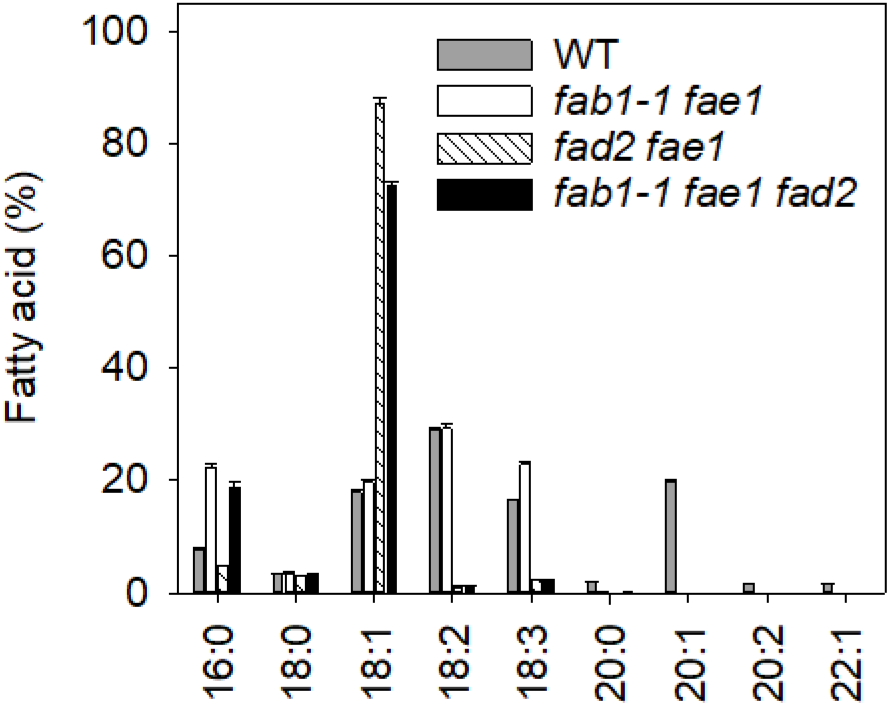
FA composition of TG from HPHO seeds. Total FA composition of TG isolated from WT, *fab1-1 fae1, fad2 fae1* and *fab1-1 fae1 fad2* seeds. Values are the mean ± SE of measurements on seeds from three plants of each genotype.

### mLPAT1 expression in HPHO seed drives 16:0 incorporation into the sn-2 position of TG

We have previously shown that expression of an ER-retargeted version of the chloroplast LPAT (ΔCTS-LPAT1 or mLPAT1) in WT Arabidopsis seeds, under the soybean glycinin-1 promoter (*ProGLY*), leads to a substantial increase in esterification of 16:0 to the sn-2 position in TG (van Erp et al., 2019). To determine what effect mLPAT1 expression has in a HPHO background, we constructed a *ProGLY:mLPAT1 fab1-1 fae1 fad2* line by crossing. When we purified TG from the seeds and performed positional analysis (van Erp et al., 2019), we found that the percentage of 16:0 at the sn-2 position (versus sn-1+3), had increased from ~3% in the *fab1-1 fad2 fae1* background to ~24% in *ProGLY:mLPAT1 fab1-1 fad2 fae1* (Fig. 3A). The total FA composition of *fab1-1 fad2 fae1* seeds was not altered greatly by mLPAT1 expression, except that there was a significant (P > 0.05) increase in 16:0 abundance, from ~19 to ~22% (Fig. 3B). We previously observed an increase in total 16:0 content when we expressed mLPAT1 in WT seeds (van Erp et al., 2019; Fig. 3B). Our data show that mLPAT1 expression in HPHO seeds allows incorporation of 16:0 into the sn-2 position of TG. However, the level of enrichment is significantly lower (P > 0.05) than in WT seeds containing *ProGLY:mLPAT1* (Fig. 3A).

**Figure 3.**
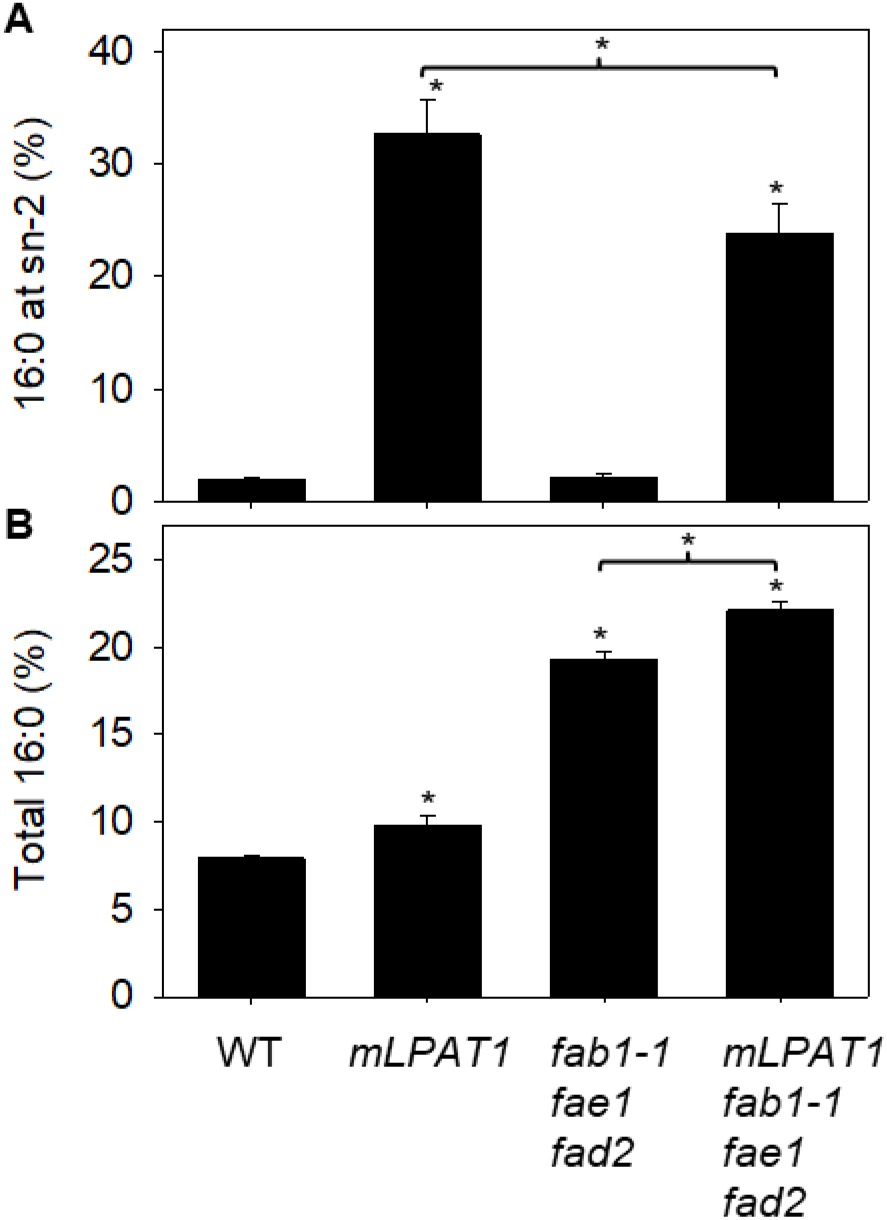
16:0 in TG from WT and HPHO seeds expressing *mLPAT1*. Percentage of 16:0 esterified to the sn-2 position (A) and 16:0 as a percentage of total FA content (B) measured in TG isolated from WT, *ProGLY:mLPAT1*, *fab1-1 fad2 fae1* and *ProGLY:mLPAT1 fab1-1 fad2 fae1* seeds. Values are the mean ± SE of measurements on seeds from three plants of each genotype. * denote values significantly (P < 0.05) different either from WT or, where marked in parenthesis, from one another (ANOVA + Tukey HSD test).

### AGPAT1 expression drives stronger 16:0 incorporation into the sn-2 position of TG

To investigate whether other ER-localised LPATs might enable Arabidopsis to incorporate more 16:0 into the sn-2 position of TG than mLPAT1, we decided to test *Homo sapiens* AGPAT1 (Agarwal et al., 2011). Human milk fat globules are secreted by lactocytes in the mammary gland epithelium (Witkowska-Zimny and Kaminska-El-Hassan, 2017). It is not known precisely which LPAT is responsible for human milk fat biosynthesis. However, *AGPAT1* is expressed in mammary epithelial cells (Lamay et al., 2013) and *in vitro* assays suggest that AGPAT1 can use 16:0-CoA as a substrate (Agarwal et al., 2011). Using transient expression in *Nicotiana benthamiana* leaves, we confirmed that AGPAT1 can localise to the ER in plant cells when it is expressed as a red fluorescent protein (RFP)-AGPAT1 fusion protein under the cauliflower mosaic virus 35S promoter (Fig. 4A). Next we transformed WT Arabidopsis plants with a *ProGLY:AGPAT1* construct in order drive strong seed-specific expression of the transgene (van Erp et al., 2019). From >40 primary transformants we selected two independent single copy T2 lines (L35 & L40) for analysis and obtained homozygous T3 seed. When we purified TG from the homozygous seed batches and carried out positional analysis, we found that the percentage of 16:0 at the sn-2 position was ~66% for L40 and ~74% for L35 (Fig. 4B). To determine what effect AGPAT1 expression has in the HPHO background we constructed a *ProGLY:AGPAT1 fab1-1 fae1 fad2* line by crossing. When we purified TG from these seeds and performed positional analysis, we found that the percentage of 16:0 at the sn-2 position was ~54% (Fig. 4B). The total 16:0 content of *ProGLY:AGPAT1 fab1-1 fae1 fad2* seeds was not significantly different (P > 0.05) from *fab1-1 fae1 fad2* (Fig. 4C). AGPAT1 expression therefore allows a higher incorporation of 16:0 into the sn-2 position of TG in WT and HPHO seeds than mLPAT1 (van Erp et al., 2019).

**Figure 4.**
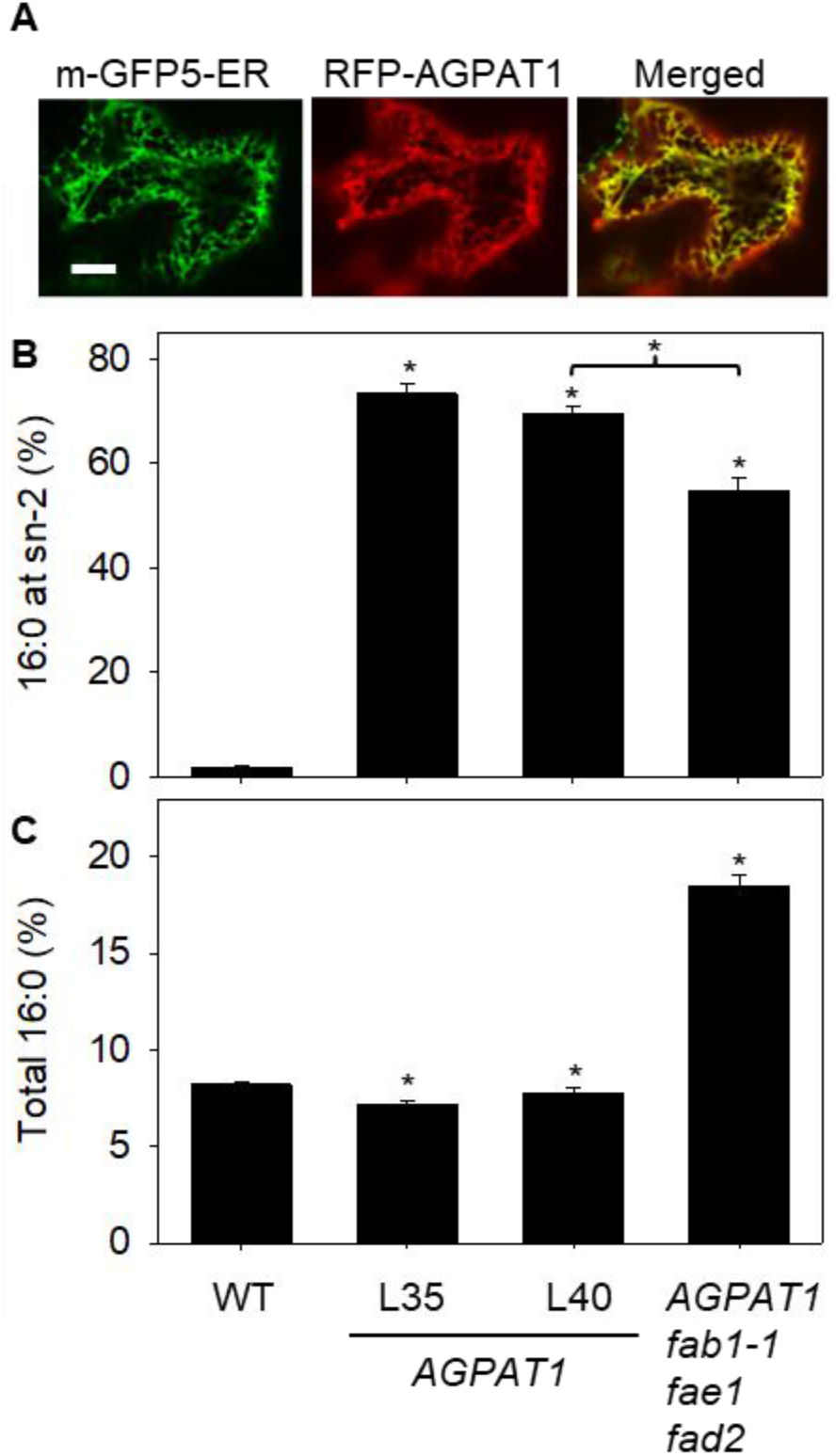
16:0 in TG from WT and HPHO seeds expressing *AGPAT1*. Laser scanning confocal microscopy image of a *N. benthamiana* epidermal cell transiently expressing RFP-AGPAT1 and m-GFP5-ER marker (A). Scale bar = 20 μm. Percentage of 16:0 esterified to the sn-2 position (B) and 16:0 as a percentage of total FA content (C) measured in TG isolated from WT, *ProGLY:AGPAT1*, *fab1-1 fad2 fae1* and *ProGLY:AGPAT1 fab1-1 fad2 fae1* seeds. Values are the mean ± SE of measurements on seeds from three plants of each genotype. * denote values significantly (P < 0.05) different either from WT or, where marked in parenthesis, from one another (ANOVA + Tukey HSD test).

### Disruption of *LPAT2* and *PDCT* enhances mLPAT1 and AGPAT1-dependent 16:0 incorporation into the sn-2 position of TG in HPHO seeds

In wild type (WT) Arabidopsis seeds we previously found that mLPAT1-dependent incorporation of 16:0 into the sn-2 position of TG could be increased by disrupting the enzymes LPAT2 and PDCT (van Erp et al., 2019). LPAT2 is the main ER-localized LPAT isoform expressed in Arabidopsis seeds (Kim et al., 2005) and therefore disruption likely reduces competition with mLPAT1 (Fig. 1) (van Erp et al., 2015). PDCT catalyses rapid DG/PC interconversion in Arabidopsis seeds (Lu et al., 2009). Although LPAT initially acylates glycerolipids at sn-2, once these acyl groups are in PC they can be removed and replaced by a deacylation-reacylation (acyl editing) cycle (Stymne and Stobart, 1984; Wang et al., 2012). Disruption of PDCT forces a more direct flux of newly made DG into TG (Fig. 1) (Bates et al., 2012). To determine whether LPAT2 and PDCT disruption affect the percentage of 16:0 esterified to the sn-2 position in TG in HPHO seeds expressing mLPAT1 or AGPAT1, we constructed *ProGLY:mLPAT1 fab1-1 fae1 fad2 lpat2-2 pdct* and *ProGLY:AGPAT1 fab1-1 fae1 fad2 lpat2-2 pdct* lines by crossing. When we purified TG from these seeds and performed positional analysis, we found that the percentage of 16:0 at the sn-2 position, was ~62% and ~84%, respectively (Fig. 5A). The total 16:0 content in *ProGLY:mLPAT1 fab1-1 fae1 fad2 lpat2-2 pdct* and *ProGLY:AGPAT1 fab1-1 fae1 fad2 lpat2-2 pdct* seeds was ~25 and ~23%, respectively (Fig. 5B). Virtually all the remaining FA was 18:1 (Fig. 2). The levels of 16:0 enrichment at sn-2 therefore compare favourably to that of human milk fat (Giuffrida et al., 2018) and a substantial proportion of the TG must also be OPO, given the total FA composition.

**Figure 5.**
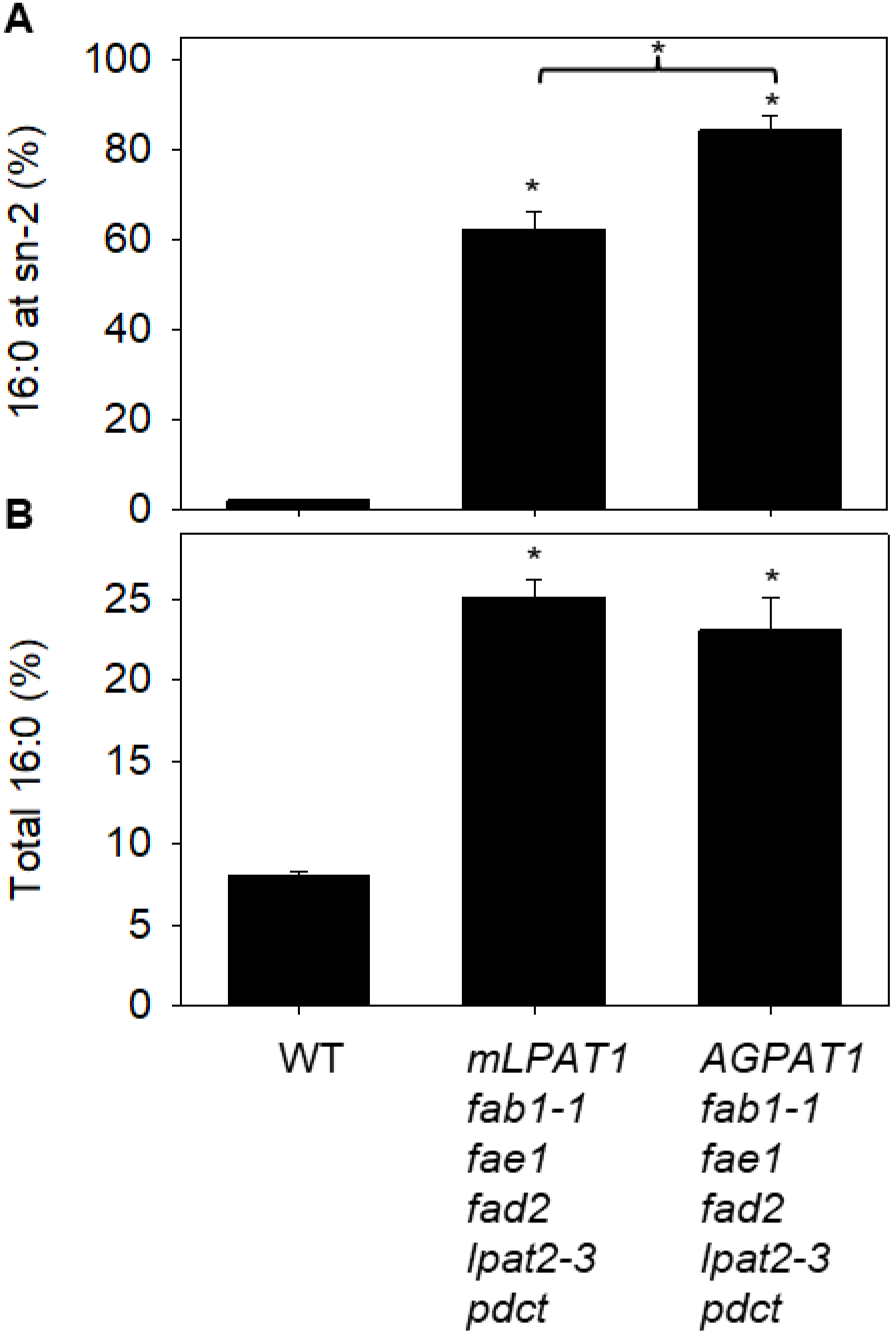
16:0 in TG from HPHO *lpat2-3 pdct* seeds expressing *mLPAT1* or *AGPAT1*. Percentage of 16:0 esterified to the sn-2 position (A) and 16:0 as a percentage of total FA content (B) measured in TG isolated from WT, *ProGLY:mLPAT1 fab1-1 fae1 fad2, lpat2-3 pdct* and *ProGLY:AGPAT1 fab1-1 fae1 fad2 lpat2-3 pdct* seeds. Values are the mean ± SE of measurements on seeds from three plants of each genotype. * denote values significantly (P < 0.05) different either from WT or, where marked in parenthesis, from one another (ANOVA + Tukey HSD test).

### HPHO seeds have reduced oil content and seed vigour but this is not compounded by redistribution of 16:0 to the sn-2 position

Modification of FA composition can reduce TG accumulation in oilseeds and can also impair seed germination and seedling establishment (Lunn et al., 2017; Bai et al., 2019). We previously found that *ProGLY:mLPAT1 lpat2-3 pdct* seeds, which have a low total 16:0 content but ~70% esterified to the sn-2 position, exhibit a reduction in TG content as a percentage of seed weight (van Erp et al., 2019). However, their germination and initial seedling growth were not significantly impaired (van Erp et al., 2019). To examine the physiological impact of 16:0 enrichment at the sn-2 position of TG in HPHO seeds, we compared seed batches from WT, *fab1-1 fae1 fad2, ProGLY:mLPAT1 fab1-1 fae1 fad2 lpat2-2 pdct* and *ProGLY:AGPAT1 fab1-1 fae1 fad2 lpat2-2 pdct* plants that had been grown together under standard laboratory conditions. We found that both seed weight and percentage oil content were significantly reduced (P > 0.05) in *fab1-1 fae1 fad2* relative to WT (Fig. 6). However, no significant difference was observed between *fab1-1 fae1 fad2* and *ProGLY:mLPAT1 fab1-1 fae1 fad2 lpat2-2 pdct* or *ProGLY:AGPAT1 fab1-1 fae1 fad2 lpat2-2 pdct* seeds (Fig. 6). These data suggest that TG biosynthetic flux is impaired in *fab1-1 fae1 fad2* seeds, but that the further genetic modifications leading to incorporation of 16:0 into the sn-2 position of TG are not detrimental in this background. These findings contrast with what we observed in WT seeds (van Erp et al., 2019). In standard laboratory conditions (20°C, 16h photoperiod), the speed of *fab1-1 fae1 fad2* seed germination (scored as radicle emergence) and seedling establishment (scored as cotyledon expansion) was significantly slower (P > 0.05) than WT (Fig. 7). However, no significant difference (P > 0.05) was observed between *fab1-1 fae1 fad2* and *ProGLY:mLPAT1 fab1-1 fae1 fad2 lpat2-2 pdct* or *ProGLY:AGPAT1 fab1-1 fae1 fad2 lpat2-2 pdct* seeds (Fig. 7). These data suggest that seed vigour is impaired in *fab1-1 fae1 fad2* seeds, but that incorporation of 16:0 into the sn-2 position of TG does not compound this phenotype.

**Figure 6.**
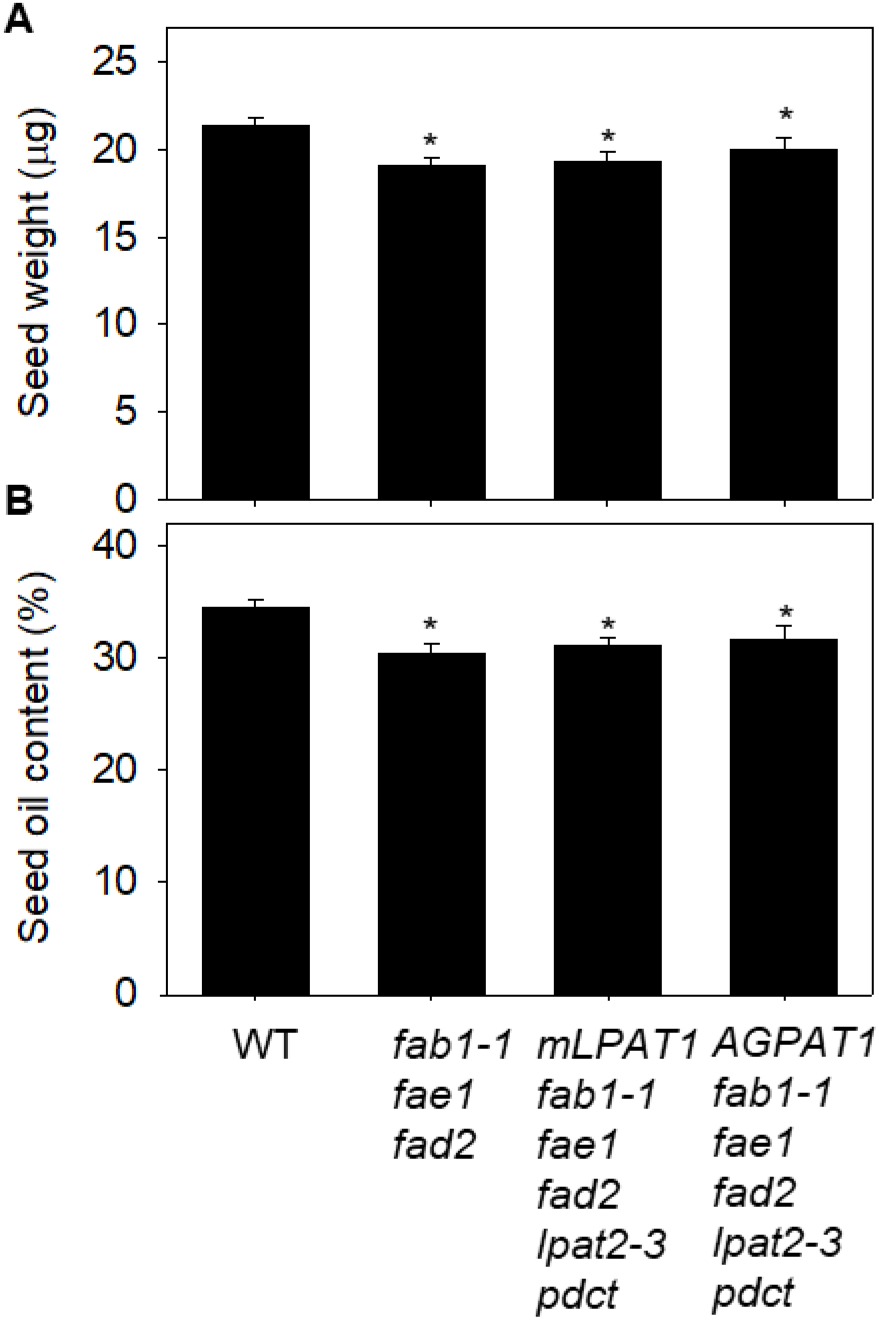
Oil content of HPHO *lpat2-3 pdct* seeds expressing *mLPAT1* or *AGPAT1*. Seed weight (A) and percentage oil content (B) of WT, *fab1-1 fae1 fad2, ProGLY:mLPAT1 fab1-1 fae1 fad2, lpat2-3 pdct* and *ProGLY:AGPAT1 fab1-1 fae1 fad2 lpat2-3 pdct* seeds. Values are the mean ± SE of measurements on seeds from three plants of each genotype. * denote values significantly (P < 0.05) different either from WT (ANOVA + Tukey HSD test).

**Figure 7.**
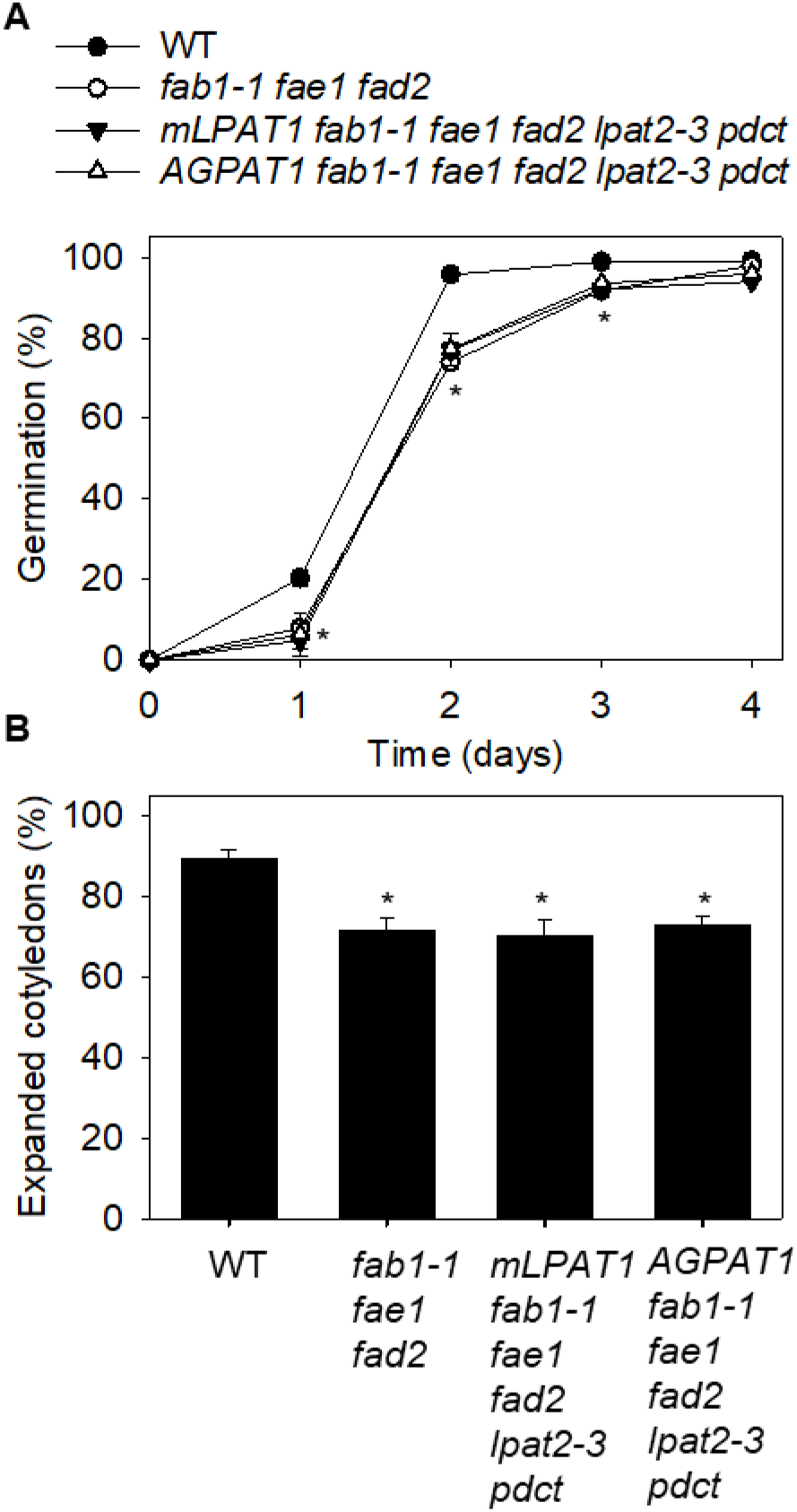
Vigour of HPHO *lpat2-3 pdct* seeds expressing *mLPAT1* or *AGPAT1*. Percentage seed germination (A) and cotyledons fully expanded by day 4 (B) of WT, *fab1-1 fae1 fad2, ProGLY:mLPAT1 fab1-1 fae1 fad2, lpat2-3 pdct* and *ProGLY:AGPAT1 fab1-1 fae1 fad2 lpat2-3 pdct* seeds. Values are the mean ± SE of measurements on seeds from three plants of each genotype. * denote values significantly (P < 0.05) different either from WT (ANOVA + Tukey HSD test).

## DISCUSSION

In this study we show that Arabidopsis seeds can be engineered to produce OPO, since the TG contains >20% 16:0 and ~70% 18:1, with >80% of the 16:0 esterified to the sn-2 position on the glycerol backbone. OPO is commonly the main TG species present in human milk (Giuffrida et al., 2018), but it is virtually absent from vegetable oils, which typically contain very little 16:0 esterified to the sn-2 position (Brockerhoff and Yurkowsk, 1966; Christie et al., 1991). The high OPO content of human milk is believed to confer nutritional benefits (Innis 2011; Béghin et al., 2018) and therefore the development of a vegetable oil that is rich in OPO might provide a new source of ingredient for infant formulas (van Erp et al., 2019).

We previously showed that lipid metabolism in Arabidopsis seeds can be engineered so that ~70% of the 16:0 present in the TG can occupy the sn-2 position (van Erp et al., 2019). However, wild type Arabidopsis seeds do not have the appropriate FA composition necessary to mimic that of human milk, which is much richer in both 16:0 and 18:1 (Wei et al., 2019). We therefore produced an Arabidopsis HPHO multi-mutant background with ~20% 16:0 and ~70% 18:1 by combining *fab1-1* (Wu et al., 1994), *fae1* (James et al., 1995) and *fad2* (Miquel and Browse, 1992) alleles. This approach is also achievable in oilseed crops since Fernández-Martínez et al., (1997) previously developed a conventional HPHO sunflower (*Helianthus annuus*) variety with seeds containing ~30% 16:0 and ~55% 18:1 by combining *fab1* and *fad2* mutant alleles (Perez-Vich et al., 2016; Schuppert et al., 2006).

The expression of an ER-retargeted version of the chloroplast LPAT (mLPAT1) in WT Arabidopsis seeds allows ~30 to 40% of the 16:0 present in the TG to occupy the sn-2 position (van Erp et al., 2019). However, when we expressed mLPAT1 in the HPHO background, we found the incorporation of 16:0 into the sn-2 position was reduced to ~20%. Disruption of LPAT2 and PDCT then lead to an increase in the percentage of 16:0 at sn-2 to ~62%. This level of 16:0 enrichment at sn-2 is also lower than we were able to achieve in WT using the same approach (van Erp et al., 2019). However, the total FA composition of HPHO is more appropriate for an infant formula ingredient and ~62% 16:0 at sn-2 still compares quite favourable with commercially-available HMFS that are produced using 1,3-regiospecific lipases (Wei et al., 2019).

Generally, in human milk fat >70% of the 16:0 is esterified to the sn-2 position of TG and this level of enrichment therefore remains the target for HMFS. The human LPAT AGPAT1 can use 16:0-CoA as a substrate (Agarwal et al., 2011) and appears to be expressed in lactocytes (Lamay et al., 2013). When we expressed AGPAT1 in WT Arabidopsis seeds we found that 60 to 70% of the 16:0 present in the TG occupied the sn-2 position. This level of enrichment is approximately twice as high as we previously achieved through expression of mLPAT1 (van Erp et al., 2019). LPAT1 is a chloroplast enzyme that uses a 16:0-ACP substrate *in vivo* (Bourgis et al., 1999), it is therefore likely to be less well adapted to function in the ER and use an acyl-CoA substrate, than AGPAT1 (Agarwal et al., 2011). When we expressed AGPAT1 in a HPHO background, incorporation of 16:0 into the sn-2 position was reduced to ~54%, but disruption of LPAT2 and PDCT then lead to an increase in the percentage of 16:0 at sn-2 to ~84%. This level of enrichment of 16:0 at sn-2 is greater than or equal to that reported in human milk fat (Giuffrida et al., 2018).

Collectively, our data suggest that when the relative amount of 16:0 (and 18:1) are increased in Arabidopsis seeds, mLPAT1 and AGPAT1 incorporate a lower percentage of 16:0 into the sn-2 position of TG. Although LPAT activity is of primary importance in determining what type of acyl group is esterified to the sn-2 position, the acyl distribution across positions is also influenced by the activities of the acyltransferases responsible for esterifying the sn-1 and sn-3 positions, and by the availability of their substrates. In Arabidopsis, GLYCEROL-3-PHOSPHATE ACYLTRANSFERASE 9 (GPAT9) (Shockey et al., 2016; Singer et al., 2016) and DIACYLGLYCEROL ACYLTRANSFERASE 1 (DGAT1) (Hobbs et al., 1999; Routaboul et al., 1999; Zou et al., 1999) are likely to be responsible for esterifying most of the acyl groups to the sn-1 and sn-3 position, respectively. These enzymes may compete with native and/or heterologous LPATs for acyl-CoA substrates. To achieve a substantially higher enrichment of 16:0 at sn-2 than 84% in Arabidopsis HPHO seeds it may be necessary to exclude 16:0 from the sn-1 and sn-3 positions by replacing GPAT9 and DGAT1 with isoforms that exhibit a stronger selectivity for unsaturated acyl groups.

Metabolic pathway engineering can often have detrimental effects on TG accumulation in oilseeds and can impair seed vigour (Lunn et al., 2017; Bai et al., 2019). We previously found that redirecting 16:0 to the sn-2 position of TG in WT Arabidopsis seeds reduced oil accumulation (van Erp et al., 2019). HPHO seeds also have a lower seed oil content than WT. However, engineering a similar shift in positional distribution of 16:0 did not lead to a further reduction. Given that WT seeds have a ~3-fold lower 16:0 content than HPHO seeds, it may be that low 16:0 availability restricts the rate of TG biosynthesis in seeds engineered to possess only 16:0-CoA LPAT activity in their ER. Conversely, the ~3-fold higher 16:0 content in HPHO seeds might conceivably restrict TG biosynthesis because the native LPATs (and other acyltransferase activities) have too little 16:0-CoA activity. HPHO seeds are also significantly impaired in seed germination and early seedling growth. However, redirecting 16:0 to the sn-2 position of TG in HPHO seeds does not appear to compound this phenotype. Poorer HPHO seed vigour may be caused by the reduction in long-chain FA unsaturation, which raises the melting temperature of the oil (Sun et al., 2014). This property is not greatly influenced by the positional distribution of the FA.

## ACKNOWLEDGMENTS

We are very grateful to Professors John Browse, John Shanklin and Ljerka Kunst for providing *pdct*, *fab1-1*, *fad2* and *fae1* single or multi-mutants. This work was funded by the UK Biotechnology and Biological Sciences Research Council through Grant BB/P012663/1.

## MATERIALS AND METHODS

### Plant material and growth conditions

The *Arabidopsis thaliana* Colombia-0 mutants *fab1-1*, *fae1*, *fad2*, *pdct* and *lpat2-3* have been described previously (Wu et al., 1994; Smith et al., 2003; Lu et al., 2009; van Erp et al., 2019). For experiments performed on media, ~50 seeds from individual plants were surface sterilized, plated on agar plates containing one-half strength Murashige and Skoog salts (Sigma-Aldrich) pH 5.7 and imbibed in the dark for 4 d at 4°C. The plates were then placed in a growth chamber set to 16-h light (photosynthetic photon flux density = 150 μmol m^−2^ s^−1^) / 8-h dark cycle at a constant temperature of 20°C. Germination (radicle emergence) and cotyledon expansion was scored visually under a dissecting stereomicroscope as described previously (van Erp et al., 2019). Individual seedlings were also transplanted to 7 cm^2^ pots containing Levington F2 compost and grown in a chamber set to a 16-h light (22 °C) / 8-h dark (16 °C) cycle, with a photosynthetic photon flux density of 250 μmol m^−2^ s^−1^. The plants were bagged individually at the onset of flowering and the seeds were harvested at maturity.

### Genotyping

Genomic DNA was isolated from leaves using the DNeasy Plant Mini Kit (Qiagen). Homozygous *lpat2-3* T-DNA insertional mutants were identified by PCR using Promega PCR Master Mix (Promega) and combinations of the gene specific and T-DNA left border primers pairs, as described previously (van Erp et al., 2019). Homozygous *fab1-1*, *fad2*, *fae1* and *pdct* mutants were identified by sequencing PCR products amplified with primer pair spanning the sites of the point mutations (Wu et al., 1994; Smith et al., 2003; Lu et al., 2009). The presence of *ProGLY:mLPAT1* and *ProGLY:AGPAT1* T-DNAs was determined by PCR using a primer pair spanning *ProGLY* and *mLPAT1* or *AGPAT1* (van Erp et al., 2019).

### Lipid analysis

Total lipids were extracted from material and TG was purified as described previously (Kelly et al., 2013). TG regiochemical analysis was performed by lipase digestion following the method described previously (van Erp et al., 2011), except that *2*-monoacylglycerols were separated by thin layer chromatography (Silica gel 60, 20 × 20 cm; Sigma-Aldrich/Merck) using hexane:diethylether:acetic acid (35:70:1.5, v/v/v) (Bates et al., 2011). Fatty acyl groups present in whole seeds and purified lipid fractions were trans-methylated and quantified by gas chromatography (GC) coupled to flame ionization detection, as described previously (van Erp et al., 2019), using a 7890A GC system fitted with DB-23 columns (30 m × 0.25 mm i.d. × 0.25 μm) (Agilent Technologies). Seed oil and moisture contents of whole seeds were measured by low-resolution time domain NMR spectroscopy using a Minispec MQ20 device (Bruker) fitted with a robotic sample-handling system (Rohasys) as described previously (van Erp et al., 2014) and the percentage oil content was normalised to 9% moisture.

### Cloning and transformation

*H. sapiens* AGPAT1 (GenBank: NP_001358367) was codon optimised for expression in Arabidopsis, synthesised by Genscript and supplied in pUC57. AGPAT1 was then amplified by PCR with KOD DNA polymerase (Merck) using primer pair 5’-CACCATGGATTTATGGCCTGGTGC-3’ & 5’-TCATCCTCCTCCACCTGG-3’. The resulting PCR product was purified with the QIAquick Gel Extraction Kit (Qiagen). The PCR product was cloned in the pENTR/D-TOPO vector (Thermo Fisher Scientific), sequenced (Fig. S1) and recombined into pK7WGR2 (Vlaams Institute for Biotechnology) using the Gateway LR Clonase II Enzyme mix (Thermo Fisher Scientific). AGPAT was cloned in the pBinGlyRed3 vector in between the soybean glycinin-1 (GLY) promoter and terminator for seed specific expression (Zhang et al., 2013). AGPAT1 was PCR-amplified from the pENTR-D-TOPO vector using KOD DNA polymerase and primer pair 5’-CGGAATTCATGGATTTATGGCCTGGTGC-3’ & 5’-GCTCTAGATCATCCTCCTCCACCTGG-3’. The PCR product was gel purified and digested with EcoRI and XbaI. The pBinGlyRed3 vector was also digested with EcoRI and XbaI, alkaline phosphatase treated (Promega), gel purified and AGPAT1 was ligated into the vector using T4 DNA ligase (NEB). Heat shock was used to transform the pK7WGR2 and pBinGlyRed3 vectors into *Agrobacterium tumefaciens* strain GV3101. Arabidopsis transformation was carried out using the floral-dip method (Clough and Bent, 1998). T1 seeds expressing the selectable marker were identified under a Leica M205 FA microscope using the DsRed filter.

### Transient expression in *N. benthamiana* and imaging

Transient expression in *N. benthamiana* leaves was carried out as described by Wood et al., (2009) using *A. tumefaciens* cultures transformed with vectors harbouring *Pro35S:RFP-AGPAT1*, *Pro35S:m-GFP5-ER* or *Pro35S:p19*. Cultures were hand-infiltrated into leaves and the inoculated plants were left for 48 h. *N. benthamiana* leaves were then mounted in water on a Zeiss LSM 780 laser scanning confocal microscope under an Apochromat 63x/1.20 W Korr M27 objective. GFP was excited at a wavelength of 488 nm and RFP at 561 nm. Filters with an emission band at 473-551 nm were used for detection.

### Statistical Analyses

All experiments were carried out using three biological replicates and the data are presented as the mean values ± standard error of the mean (SE). For statistical analysis we either used one-way analysis of variance (ANOVA) with post-hoc Tukey HSD (Honestly Significant Difference) tests, or two-tailed Student’s t-tests.

## Accession Numbers

Sequence data from this article can be found in the GenBank/EMBL data libraries under accession numbers: NP_001358367 (AGPAT1), AF111161 (LPAT1), At1g74960 (FAB1), At3g12120 (FAD2), At4g34520 (FAE1), At3g57650 (LPAT2), At3g15820 (PDCT).

## Supplemental Data

**Supplemental Figure S1.**
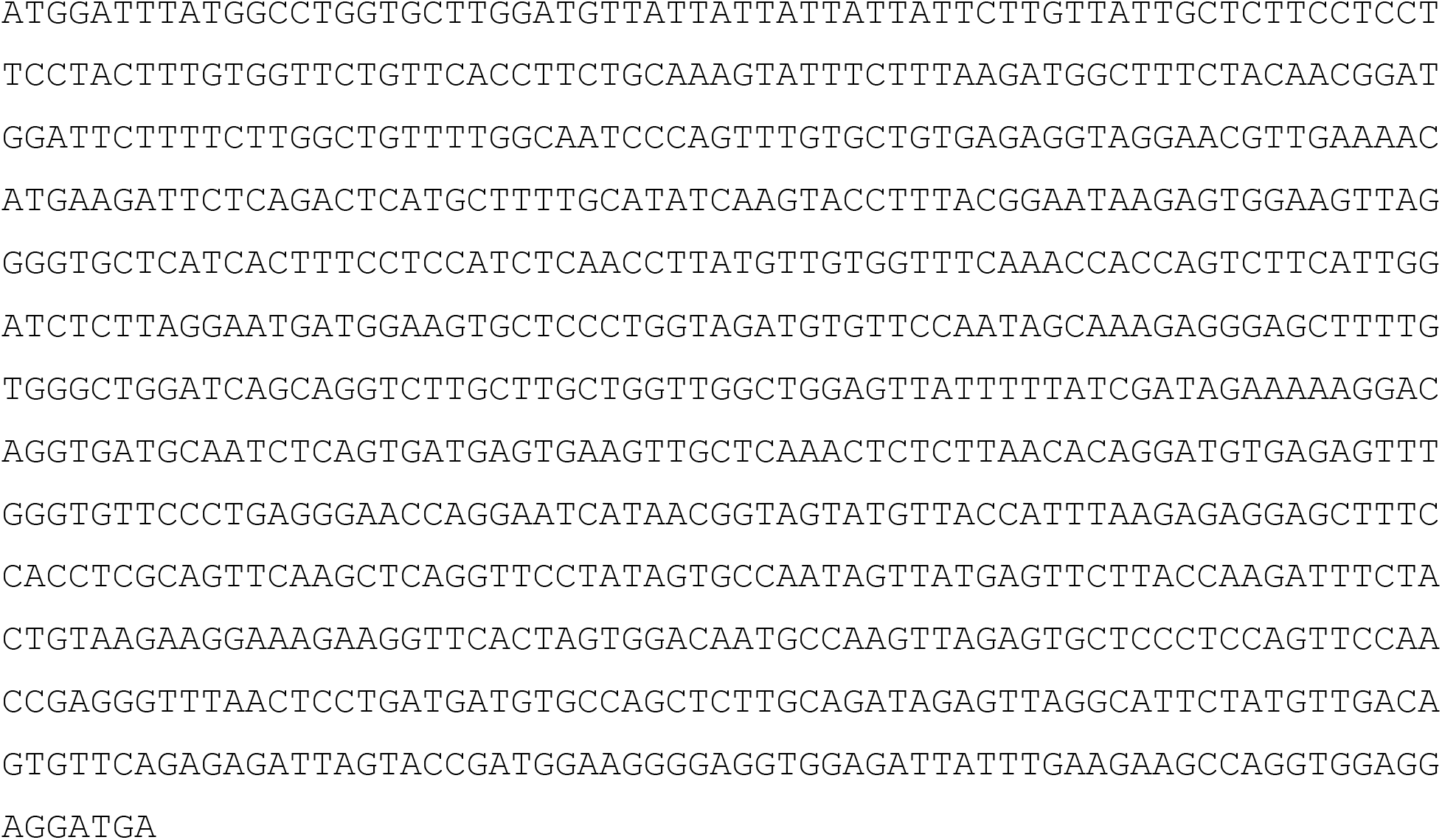
Codon optimised AGPAT1 sequence.

## Notes

**Footnotes:** This work was supported by the UK Biotechnology and Biological Sciences Research Council through an Institute Strategic Program Grant and BB/E022197/1.

### Competing Interest Statement

The authors have declared no competing interest.

